# Domain classification of archaeal proteomes reveals conserved fold repertoire

**DOI:** 10.64898/2026.04.01.715805

**Authors:** R. Dustin Schaeffer, Jimin Pei, Rui Guo, Jing Zhang, Kirill Medvedev, Qian Cong, Nick Grishin

## Abstract

Archaea represent one of the three domains of cellular life and yet account for fewer than 1% of experimentally determined protein structures, leaving the extent of their structural novelty unknown. Here we present a systematic domain-level classification of 124,075 proteins from 65 archaeal classes spanning 21 phyla and all major lineages, using both AFDB and newly predicted AlphaFold3 structures classified against the Evolutionary Classification of protein Domains (ECOD). We assigned 204,758 domains, of which 76.8% received high-confidence classifications, spanning 987 ECOD X-groups; 40% of known structural diversity within a single domain of life. Clustering by Foldseek recovered structural relationships for 63% of domains that are singletons by sequence comparison. To characterize the 21% of proteins lacking high-confidence classification, we applied successive filters for structure prediction confidence, protein length, and structural cluster context, reducing 8,452 domain-free proteins to a small number of well-folded structural orphans (less than 0.1% of the dataset). The unclassified fraction is dominated by sub-threshold matches to known folds (14% of all proteins) and low-confidence structure predictions (5%), not by novel structures. These results demonstrate that the protein fold repertoire at the single-domain level is broadly conserved across the deepest phylogenetic distances in cellular life, and that the gap between archaeal and well-characterized proteomes reflects classification sensitivity for divergent sequences rather than unexplored structural diversity.

## Introduction

Classifying the universe of protein domains requires structural data drawn broadly from across the tree of life. The Evolutionary Classification of protein Domains (ECOD) organizes domains into a hierarchy in which homologous superfamilies (H-groups) contain domains with common ancestry, while X-groups gather H-groups whose evolutionary relationship to each other has some evidence but remains uncertain [1, 2]. ECOD currently classifies over 2.7M protein domains from both experimental and predicted structures [3], but the taxonomic distribution of its source data is heavily skewed towards bacteria and eukaryotes. When structural data are drawn predominantly from a subset of organisms, relationships that exist in nature may go undetected, not because homology is absent, but because the intermediates needed to reveal it have not been sampled.

Archaea represent one of the most extreme cases of this sampling asymmetry. Though recognized as a primary domain of cellular life, whether as the third domain in the classical Woese framework or as the clade from which eukaryotes emerged in two domain models[4, 5], archaea account for approximately 2.5% of experimentally determined structures in the Protein Data Bank (PDB) and 2.7% of sequences in the AlphaFold Protein Structure Database (AFDB) [6]. This underrepresentation is not a minor gap: archaea span enormous phylogenetic and ecological diversity, from the reduced-genome DPANN lineages to the eukaryote-related Asgard archaea[7], and occupy environments ranging from hydrothermal vents and hypersaline lakes to temperate soils and ocean waters [8–13] and the human gut microbiome [14]. Their proteins have evolved under selective pressures that differ from, and in many cases are more varied than, those shaping bacterial and eukaryotic proteomes, potentially driving exploration of regions of sequence and structure space that other lineages have not sampled. Prior phylogenomic analyses of domain structures across proteomes have noted archaea’s minimal structural repertoire[15], but these studies were limited to experimentally determined structures from cultivated organisms. Recent analysis of nearly 3,000 archaeal genomes found that even standard bioinformatic pipelines leave approximately 42% of archaeal genes uncharacterized, with long stretches of genes lacking even Pfam domain annotation [16], underscoring the gap between archaeal diversity and current classification tools. If novel protein folds remain to be discovered, archaeal proteomes are among the most likely places to find them.

Advances in protein structure prediction now make systematic surveys of underrepresented lineages feasible. AlphaFold2 and AlphaFold3 produce predicted structures with backbone accuracy approaching experimental methods for most globular proteins, though with reduced reliability for side chains and regions lacking homologous templates [17, 18]. Automated domain classification pipelines such as DPAM (Domain Parser for AlphaFold Models) can assign predicted structures to ECOD domains through iterative sequence and structural comparison [19]. These tools have been applied to proteome-scale datasets including 48 model organism and pathogen proteomes from the AlphaFold Database [20, 21], but no prior study has applied domain-level structural classification to archaeal proteomes at comparable scale. Meanwhile, structure-based comparison methods such as Foldseek detect remote homology well beyond the reach of sequence comparison [22], enabling both the recovery of relationships invisible to sequence methods and the identification of proteins with no detectable similarity to any known fold.

Here we present a systematic structural survey of 124,075 proteins from 65 archaeal classes spanning 21 phyla and all major archaeal lineages. Using existing structures from the AFDB and de novo AlphaFold3 predictions, we classified 204,758 protein domains against ECOD and annotated them against Pfam, producing a comprehensive archaeal domain classification. Structural clustering by Foldseek finds significant similarity for 63% of domains that are singletons by sequence comparison. Most significantly, we find that existing domain templates classify approximately 80% of archaeal proteins with high-confidence. Furthermore, systematic characterization of the unclassified fraction reveals that it is dominated by classification sensitivity limits and structure prediction quality rather than by novel structures. These results indicate that the protein fold repertoire at the single-domain level is broadly conserved across cellular life, and point toward family-level expansion, classifier sensitivity, and domain architecture diversity as the productive frontiers for structural classification.

## Results

### Dataset composition and structure prediction

The archaeal proteome dataset comprises 124,075 proteins spanning 65 Genome Taxonomy Database (GTDB) classes, 21 phyla, and 6 archaeal major groups (**Fig 1A**). Proteins were drawn from three sources: 71,866 from the AFDB, 29,326 from Prodigal gene predictions (structures predicted *de novo* by AF3), and 22,883 from UniParc sequence matches (also predicted by AF3). The six groups ranged from 6,543 (Deep-branching) to 32,066 proteins (Euryarchaeota-related), with DPANN (7,887), Thermoplasmatales (22,422), the TACK superphylum (23,641), and Asgard archaea (31,516) in between. The relative contribution of each data source varied by lineage: AFDB provided the majority of Euryarchaeota-related (83.4%) and TACK (78.9%) proteins, whereas Asgard archaea and Thermoplasmatales required predominantly new structure predictions, with AFDB contributing only 35.1% and 26.3% of proteins, respectively.

**Figure 1.**
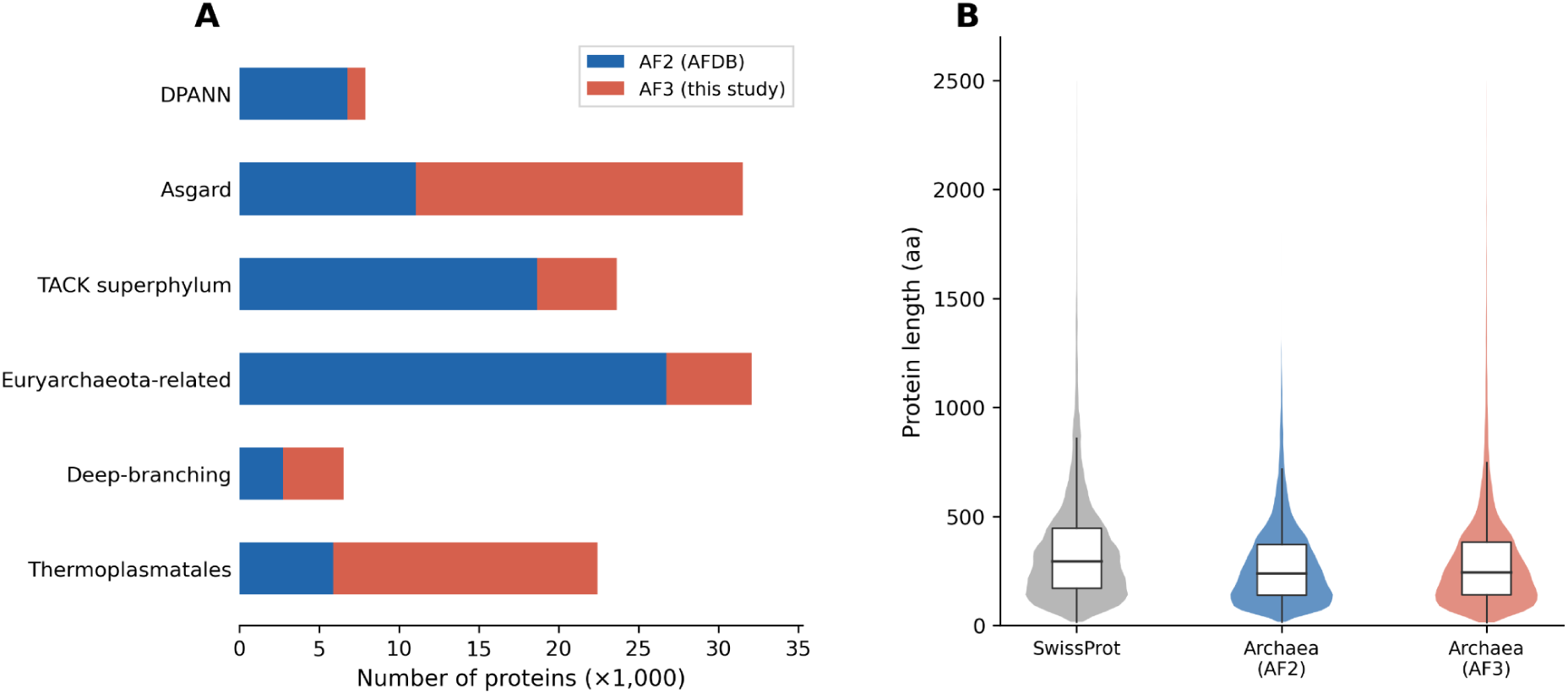
Overview of the archaeal proteome dataset. **(A)** Dataset composition by major archaeal group and structure prediction method. Stacked horizontal bars show the number of proteins with structures from the AlphaFold Database (AF2; blue) versus those predicted de novo in this study (AF3; red). Euryarchaeota-related and TACK superphylum lineages are well-represented in AFDB, whereas Asgard archaea and Thermoplasmatales required predominantly new structure predictions. The dataset totals 124,075 proteins across 65 classes, 21 phyla, and 6 major groups. **(B)** Protein length distributions for Swiss-Prot (n = 542,378; median 295 aa), AF2 (n = 71,863; median 240 aa), and AF3 (n = 52,360; median 245 aa). Violin plots show kernel density estimates; embedded box plots indicate the interquartile range and median. Archaeal proteins are shorter than the Swiss-Prot reference, consistent with the compact genomes characteristic of prokaryotic organisms. The two archaeal subsets show similar length distributions, indicating that prediction method does not introduce a systematic size bias. Proteins longer than 2,500 aa are excluded for display.

Archaeal proteins are shorter than those in Swiss-Prot, with a median length of 242 amino acids compared to 295 for Swiss-Prot (n = 542,378) (**Fig 1B**). The AF2 (AFDB) and AF3 subsets showed similar length distributions (median 240 vs. 245 amino acids), indicating that data source did not introduce a systematic size bias. AlphaFold2 and AlphaFold3 structures achieved comparable prediction confidence: AF2 structures had a median per-residue pLDDT of 89.3 compared to 86.2 for AF3, with similar residue coverage and domain density (**S1 Fig**). The modest difference in DPAM judge distributions, 79.9% high-confidence for AF2 versus 72.4% for AF3, concentrated in the low-confidence category, does not compromise downstream analyses, which apply uniform quality filters. Of the 124,075 proteins, 116,275 (93.7%) have structure quality metrics, with an overall median pLDDT of 85.7.

### ECOD classification landscape

DPAM assigned 204,758 domains across 115,623 proteins. Of these, 157,287 (76.8%) received high-confidence classifications (“well assigned”), representing strong matches to ECOD domain templates. The most abundant structural families are broadly distributed across archaeal lineages **(Fig 2A)**. The top 15 H-groups, including HTH domains (H-group 101.1), P-loop NTPases (H-group 15.1), and Rossmann-like folds (H-group 2003.1), account for 62,439 of 157,287 high-confidence domain assignments (39.7%), with representatives present in all six major archaeal groups. Archaeal domains span a substantial fraction of known structural diversity (**Fig 2B**): 987 of 2,457 ECOD X-groups (40.2%), 1,583 of 3,716 H-groups (42.6%), and 1,751 of 3,953 T-groups (44.3%). For comparison, Swiss-Prot (representing all three domains of life) covers 1,874 X-groups, 2,851 H-groups, and 3,060 T-groups. Single-domain proteins predominate (63,473 of 115,623; 54.9%), followed by two-domain proteins (32,370; 28.0%), with progressively fewer multi-domain architectures up to a maximum of 20 domains per protein **(Fig 2C)**.

**Figure 2.**
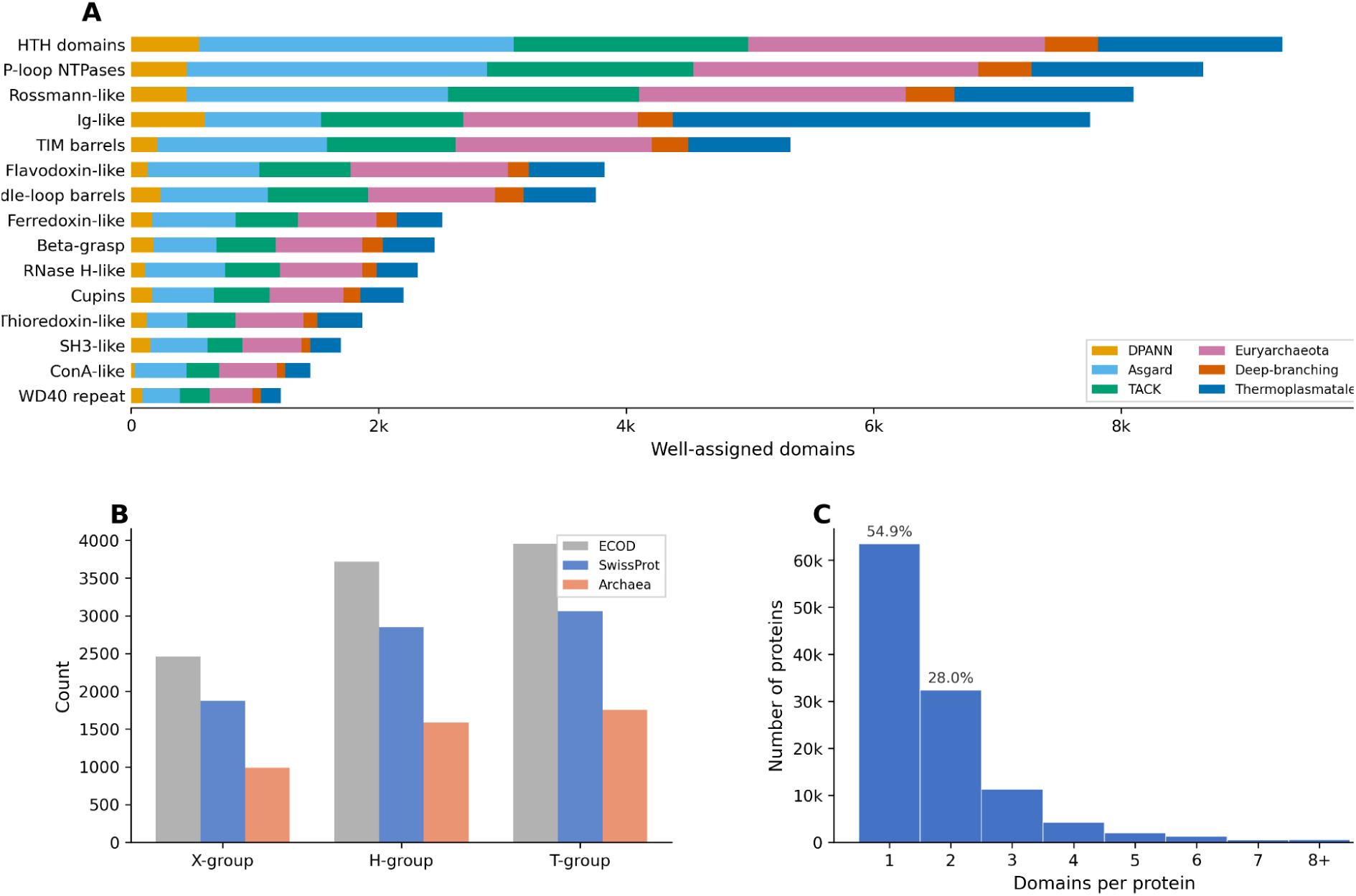
DPAM domain classification landscape across the archaeal proteome. **(A)** Top 15 ECOD H-groups (homologous superfamilies) by domain count, restricted to well-assigned domains. Stacked bars show contributions from each major archaeal lineage. HTH domains, P-loop NTPases, and Rossmann-like folds are the most abundant structural families, with broad distribution across all archaeal groups. The top 15 H-groups account for 62,439 of 157,287 well-assigned domains (39.7%). **(B)** ECOD hierarchy coverage comparison. Archaeal domains span 987 X-groups (40% of 2,457 in ECOD), 1,583 H-groups (43% of 3,716), and 1,751 T-groups (44% of 3,953). Swiss-Prot covers 1,874 X-groups, 2,851 H-groups, and 3,060 T-groups. A single domain of life thus captures over 40% of known structural diversity. **(C)** Distribution of domains per protein for all 115,623 proteins with domain assignments (median = 1). Single-domain proteins predominate (54.9%), with 28.0% having two domains and progressively fewer multi-domain architectures.

To assess the global impact of expanded archaeal sampling on ECOD taxonomic occupancy, we compared the distribution of H-groups across the three superkingdoms before and after incorporation of the archaeal proteome dataset (**Fig 3**). The most prominent change is a marked expansion of the universal set: H-groups present in Eukaryota, Bacteria, and Archaea increased from 746 to 1,326. In parallel, the number of H-groups assigned only to Eukaryota and Bacteria decreased from 1,329 to 879, while eukaryote-only and bacteria-only categories also contracted. By contrast, archaea-only H-groups increased only marginally, from 34 to 39. These shifts indicate that many H-groups previously interpreted as absent from Archaea were in fact unsampled, and that archaeal domains occupy a substantially broader fraction of the shared cellular fold repertoire than suggested by earlier datasets.

**Figure 3.**
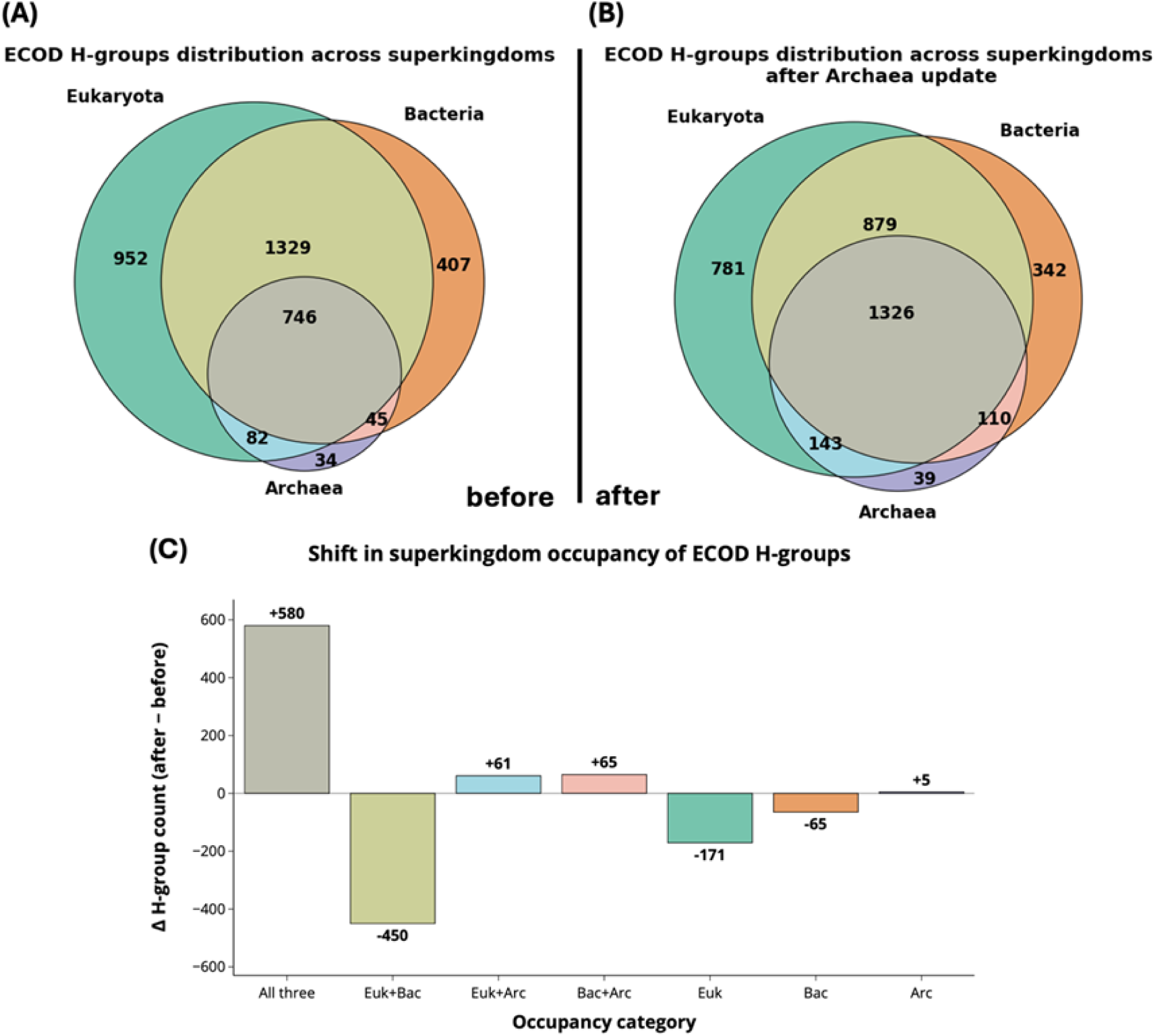
Expanded archaeal sampling redistributes ECOD H-groups from apparently lineage-restricted categories into shared and universal sets. (A,B) Venn diagrams showing the distribution of ECOD H-groups across Eukaryota (E), Bacteria (B), and Archaea (A) in the full ECOD database before (A) and after (B) incorporation of the expanded archaeal proteome dataset. (C) Bar plot showing the change in H-group counts for each superkingdom occupancy category after archaeal expansion.

To place this shift in context, we asked how well a single archaeal genome captures archaeal structural diversity. *Methanocaldococcus jannaschii* was selected for this comparison because it is the sole archaeal representative among the 48 model proteomes provided by the AlphaFold Protein Structure Database. When *M. jannaschii* alone is used to represent Archaea, 13 H-groups are classified as archaea-specific and 542 as universal. Expanding to all archaeal proteomes increases these values to 34 and 746, respectively, in the pre-expansion ECOD dataset, and further to 39 and 1,326 after archaeal expansion. Thus, a single archaeon recovers a substantial portion of deeply conserved folds but underestimates both the breadth of archaeal specialization and the extent of fold sharing across superkingdoms.

Notably, even with broad archaeal sampling, the number of archaea-exclusive H-groups remains small. The few additional archaeal-specific folds identified after expansion are predominantly associated with niche or lineage-restricted functions, including CRISPR defense systems, mobile-element or viral-related proteins, and uncharacterized families, rather than core metabolic or informational processes. At the same time, many H-groups previously assigned to eukaryotes, bacteria, or their intersection are reassigned to shared categories, particularly the universal set.

These observations show that expanded archaeal sampling primarily reveals previously unsampled members of existing fold families rather than introducing substantial new fold diversity. Archaeal proteomes therefore appear to be constructed largely from a common cellular fold repertoire, with lineage-specific innovation arising mainly through specialization and diversification within a limited set of structural frameworks.

### Domain repertoire across archaeal lineages

The six major archaeal groups share a common domain repertoire that varies in size but not kind. H-group counts vary from 797 (DPANN) to 1,228 (Euryarchaeota-related), largely tracking genome size; 96% of DPANN H-groups are shared with other archaea. The 31 DPANN-unique H-groups are predominantly singletons (27 of 31) drawn from families whose ECOD representation is otherwise bacterial or eukaryotic (e.g., chitin-binding domains, Kazal-type protease inhibitors), consistent with either horizontal gene transfer or classification artifacts in highly-reduced genomes, though individual cases have not been systematically verified. Of 592 H-groups significantly enriched in archaea relative to other cellular organisms (FDR < 0.01), 70% are universal (present in all three domains of life) indicating that archaeal enrichment reflects expansion within shared families rather than innovation of new ones. The 43 enriched H-groups shared exclusively between archaea and eukaryotes but absent from bacteria are predominantly information-processing families: RNA polymerase subunits, tRNA splicing endonucleases, DNA replication initiation factors, and translation elongation factors. This structural confirmation of the archaeal-eukaryotic informational core [23] demonstrates that the shared heritage is visible at the domain level across all major archaeal lineages, not only in cultivated model organisms.

Pan-archaeal scope also recontextualizes recent reports of eukaryotic signature proteins (ESPs) in Asgard archaea. Our classification recovers known ESP families in Asgard lineages, consistent with recent structural analyses [7, 24] and characterizations of defense system repertoires shared between Asgard archaea and eukaryotes[25]. However, several protein families reported as Asgard-eukaryote connections, including the major vault protein (MVP; H-group 3529.1), classified in ECOD since 2014, are present with high-confidence across all six major archaeal groups, with Asgard accounting for only 16% of MVP domain assignments. The distinction between Asgard-specific and pan-archaeal distribution has implications that are beyond the scope of this study but merit investigation with expanded sampling.

### Annotation landscape and the unclassified fraction of domains

We assessed the completeness and reliability of ECOD classifications by comparison with Pfam annotations. Pfam concordance varied systematically by DPAM judge category **(Fig 4A)**: high-confidence assignments showed a 73.8% Pfam mapping rate (116,115 of 157,287), declining through partial domain (36.7%), low confidence (25.1%), and simple topology (12.1%). The declining Pfam rate across judge categories is consistent with DPAM’s confidence rankings reflecting genuine differences in sequence divergence from characterized families, not arbitrary thresholds.

**Figure 4.**
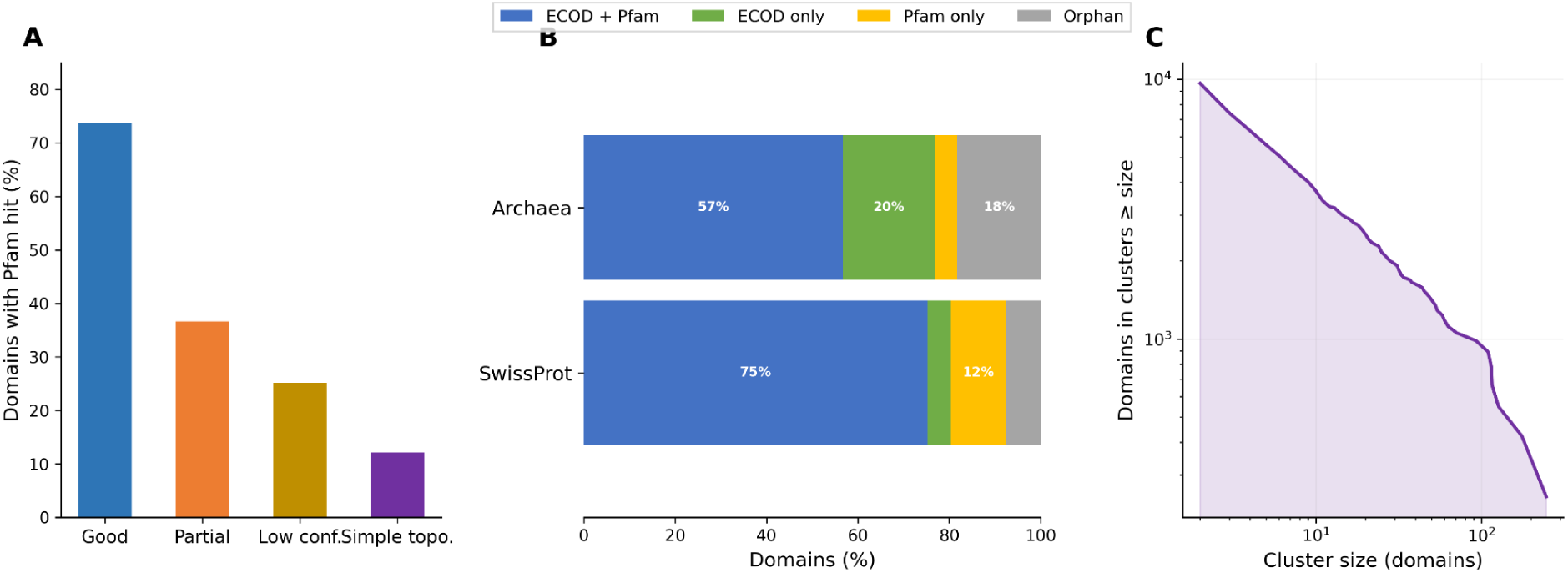
Pfam concordance and novel domain landscape. **(A)** Pfam mapping rate by DPAM classification judge. Well-assigned domains show the highest Pfam concordance (73.8%, 116,115 of 157,287), followed by partial (36.7%), low confidence (25.1%), and simple topology (12.1%). The declining Pfam rate across judge categories is consistent with increasing sequence divergence from characterized families. **(B)** Domain annotation landscape comparing archaeal and Swiss-Prot proteomes. Four categories are shown: ECOD + Pfam (both structural and sequence classification), ECOD only (structural match without Pfam), Pfam only (sequence match without high-confidence ECOD), and Orphan (neither). Archaea show 56.7% ECOD + Pfam, 20.1% ECOD only, 4.9% Pfam only, and 18.2% orphan domains. Swiss-Prot shows 75.2% ECOD + Pfam and only 7.7% orphan domains. The higher orphan fraction in archaea reflects their underrepresentation in current classification databases. **(C)** Novel domain cluster size distribution (Tier 2 orphan domains: no Pfam hit, sub-threshold DPAM assignment). Foldseek structural clustering of 37,338 orphan domains yields 29,817 clusters, of which 2,097 are multi-member. The CCDF shows the cumulative number of domains in clusters at or above each size, with the largest clusters containing hundreds of structurally similar orphan domains, candidates for novel or highly divergent domain families.

A four-way annotation landscape (**Fig 4B**) reveals that 56.7% of archaeal domains carry both a high-confidence ECOD classification and a Pfam hit, 20.1% carry ECOD only, 4.9% carry Pfam only, and 18.2% lack both (“orphan” domains). In our classification of Swiss-Prot proteins [26], orphans constitute only 7.7% of domains. The elevated orphan fraction in archaea, more than double the Swiss-Prot rate, raises a question that motivates the remainder of this analysis: does this gap represent genuine structural novelty among archaeal proteins, or is it a consequence of classifying deeply divergent sequences against a reference database built predominantly from bacterial and eukaryotic structures?

To begin addressing this question, we examined the 25,965 proteins that lack any high-confidence domain assignment (**Fig 4C**). Of these, 67.5% have at least one DPAM domain hit below the high-confidence threshold; they are sub-threshold matches to known folds, not structurally uncharacterized. The remaining 32.5% lack any DPAM domain assignment whatsoever. These domain-free proteins are characterized in detail below; as we show, the majority are explained by low-confidence structure predictions and short protein length rather than by novel folds.

### Structural clustering reveals relationships beyond sequence detection

To explore relationships beyond the reach of sequence-based classification, we performed structural clustering using Foldseek at four levels: protein sequence clusters (PSC), protein structural clusters (PXC), domain sequence clusters (DSC), and domain structural clusters (DXC). Sequence-based clustering identified a high proportion of singletons: 53.2% at the protein level and 51.7% at the domain level (**Fig 5A**). Structure-based clustering substantially reduced these fractions to 15.2% and 20.2%, respectively. At the domain level, 66,399 sequence singletons were rescued into multi-member structural clusters, a 63% recovery rate. The greater sensitivity of structural clustering is evident in cluster size distributions (**Fig 5B**): structural clusters consistently show heavier tails than their sequence counterparts on log-log plots. Many DXC clusters unify members from multiple DSC clusters (**Fig 5C**), with 9,800 DXC clusters bridging two or more DSC clusters, indicating that archaeal protein families harbor far more internal diversity than sequence comparison alone can capture. Clustering parameters (E-value 0.001, 50% coverage, TM-align mode) were chosen for sensitivity in detecting remote relationships; structural similarity at these thresholds indicates shared topology but does not by itself establish common evolutionary origin, particularly for abundant folds such as Rossmann-like and TIM barrel domains where convergence is well documented.

**Figure 5.**
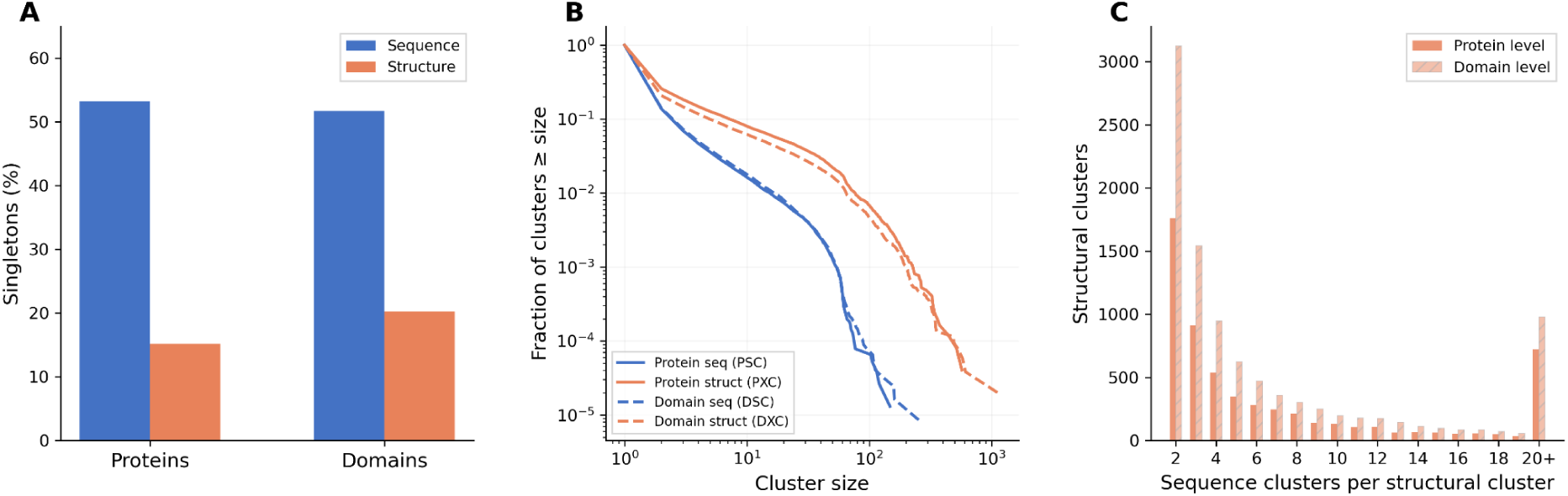
Structural clustering reveals relationships beyond sequence detection. **(A)** Singleton fractions for sequence-based versus structure-based clustering at both protein and domain levels. Structure-based clustering recovers 63% of domain-level sequence singletons into multi-member clusters. **(B)** Complementary cumulative distribution functions (CCDFs) of cluster sizes across all four clustering dimensions on a log-log scale. Structural clusters show heavier tails than sequence clusters. **(C)** Number of distinct sequence clusters unified within each structural cluster. At the domain level, 9,800 structural clusters contain members from two or more sequence clusters.

### Characterizing the structural dark matter

We focused on the 8,452 proteins lacking any DPAM domain assignment and applied successive filters to identify genuinely novel structures (**Fig 6A**). The largest filter is structure quality: 74.0% have a mean pLDDT below 70, indicating disordered regions or poor structure predictions unlikely to harbor classifiable domains. The dramatic separation in pLDDT distributions between classified proteins (median 89.6), sub-threshold proteins (median 76.7), and domain-free proteins (median 62.6) confirms that unclassifiable proteins are predominantly those for which AlphaFold did not produce confident structures (**Fig 6B**).

**Figure 6.**
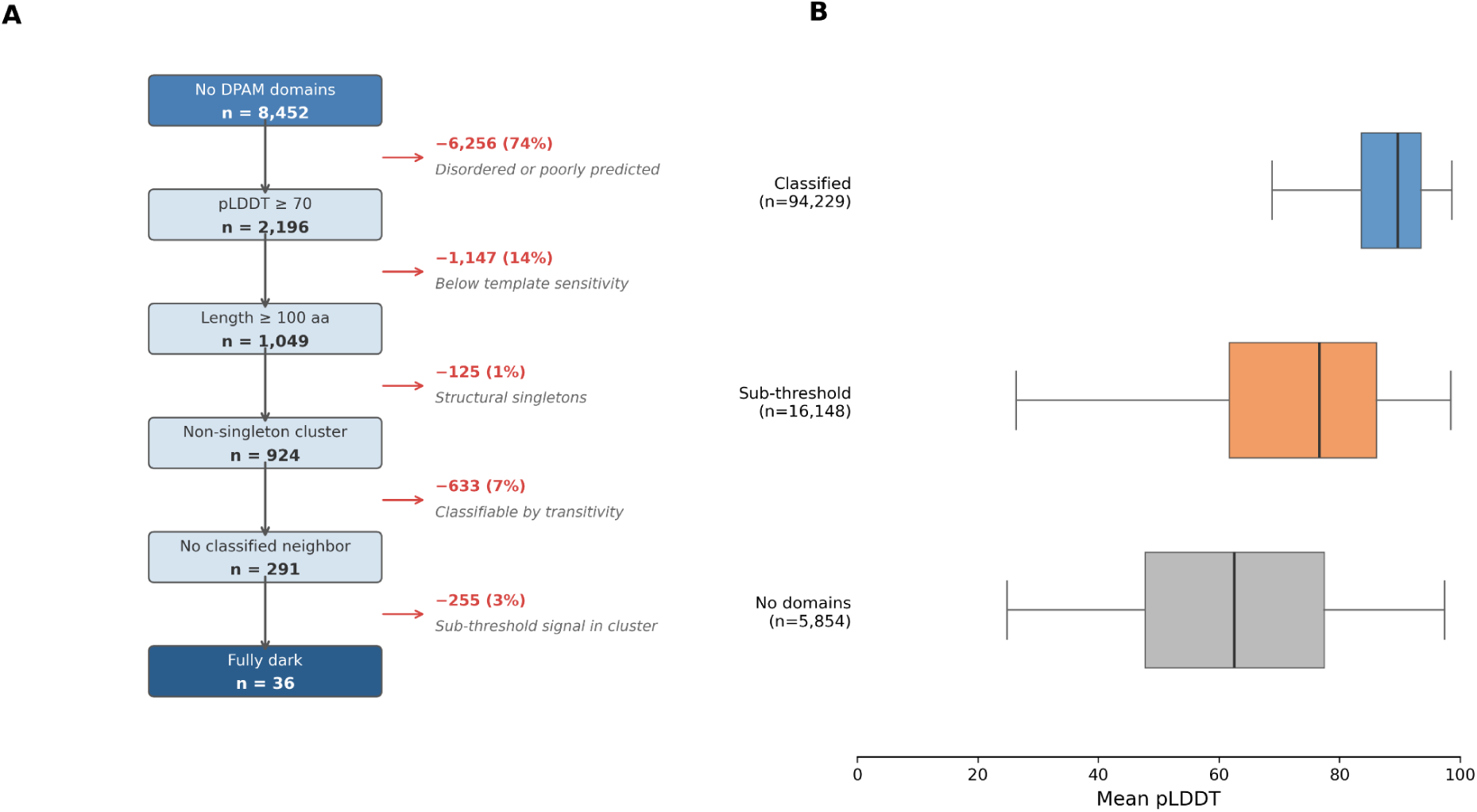
Systematic characterization of the unclassified fraction reveals that genuine structural novelty is vanishingly rare. **(A)** Dark matter filtering pipeline. Of 8,452 proteins lacking any DPAM domain assignment, successive filters remove those with low-confidence structures (pLDDT < 70; 74%), lengths below template sensitivity (< 100 aa; 14%), structural singletons (1%), membership in clusters containing classified members (7%), and membership in clusters with sub-threshold domain hits (3%). The residual, 36 proteins in 20 fully dark clusters, represents the irreducible core of potentially novel folds, constituting 0.03% of the 124,075-protein dataset. Of these 20 clusters, 18 are pairs (size 2); only PXC_023126 (10 members, 8 lineages; maps to Pfam DUF2769, present in ECOD but missed by DPAM) and PXC_008059 (10 members, 10 lineages spanning both Asgardarchaeota and Thermoproteota; 80% AFDB-source; no Pfam or InterPro annotation) are substantial. PXC_008059 represents the strongest candidate for a genuinely novel fold. **(B)** pLDDT distributions by classification category. Proteins with high-confidence ECOD classification (blue; median 89.6, n = 94,229) are strongly enriched for well-folded structures, while proteins lacking any domain assignment (gray; median 62.6, n = 5,854) are predominantly disordered or poorly predicted. Sub-threshold proteins (orange; median 76.7, n = 16,148) occupy an intermediate range. Domain rescue by structural clustering (not shown): of 10,148 sub-threshold domains in structurally consistent clusters, 82% agree with the X-group consensus of their classified cluster neighbors, rescuing 3,463 proteins from unclassified to classifiable status.

The unclassified fraction distributes unevenly across lineages. Classification rates range from 82.5% (Euryarchaeota-related) to 74.7% (DPANN). DPANN has the highest no-domain rate (9.0%) but the lowest median pLDDT among domain-free proteins (53.9), confirming that its unclassified fraction reflects structure prediction quality in small, fragmentary genomes rather than novel folds. DPANN contributes only 6 of the final well-folded dark candidates after filtering. Asgard contributes the most (45% of candidates), and Prodigal-source proteins are overrepresented relative to their dataset share, consistent with gene prediction artifacts inflating apparent novelty. The concentration of residual dark matter in Asgard (the lineage with the most distinctive cell biology) is the one pattern consistent with genuine structural novelty rather than technical artifact.

Of the 2,196 well-folded domain-free proteins, 1,147 are shorter than 100 amino acids – below the effective sensitivity of template-based domain classification – and 125 are singletons in structural clustering with no independent structural support. This leaves 924 well-folded, reasonably sized proteins in non-singleton Foldseek clusters. Most of these cluster with proteins that do carry ECOD domains: 633 are in clusters containing at least one high-confidence domain member (i.e., they are classifiable by transitivity), and 255 are in clusters with sub-threshold domain members (i.e., also have partial classification signal). The residual consists of a small number of proteins distributed across fully dark clusters, clusters where no member has any ECOD domain assignment at any confidence level. These clusters, which include a mix of data provenance and phylogenetic breadths, are candidates for genuine structural novelty and are currently under expert curation to distinguish bona fide novel folds from gene prediction artifacts and extreme classification sensitivity failures.

We also assessed the reliability of DPAM classifications using structural clustering as a validation (**Fig 5C**). Of 3,662 DXC clusters containing both high-confidence and sub-threshold domain members, 91.4% show consistent X-group consensus among their high-confidence members. Among the 10,148 sub-threshold domains in these consistent clusters, 81.5% carry a prior X-group assignment that agrees with the cluster consensus, 16.7% disagree, and 1.7% receive a new X-group assignment that they previously lacked. The disagreements are concentrated at multi-domain boundaries involving HTH motifs, where domain parsing ambiguity rather than incorrect fold assignment is the likely cause. We restricted rescue to clusters with unanimous high-confidence consensus, a conservative design reflecting the fact that structural cluster membership alone is not sufficient evidence for homology assignment, particularly for common folds where structurally similar domains may belong to different evolutionary lineages. Under this criterion, 3,463 proteins are reclassifiable, bringing the effective classification rate from 79.1% to approximately 81.9%.

Manual analysis of genuinely dark archaeal protein domains revealed several notable cases. These domains were classified by their overall secondary-structure content, three-dimensional architecture, and size. Among them, seven small cysteine-rich domains are likely to bind metals such as zinc (**Fig 7A**). Two of these (EPP00014805 and EPP00002661) contain eight conserved cysteines each and may therefore coordinate multiple metal ions (**Fig 7A**, leftmost two structures). Together with another small domain (third structure from the left in **Fig 7A**, ECOD: EPP00018990), they may represent previously unrecognized metal-binding folds. In contrast, four other small cysteine-rich domains show structural similarity to known zinc fingers. One domain (EPP00027814) appears to be a remote homolog of the zf-HC3 family in Pfam (PF16827), whereas the other three resemble treble-clef zinc fingers [27], which are defined by a zinc knuckle (a beta-hairpin containing two zinc-binding residues) followed by an alpha-helix whose N-terminus contributes two additional zinc-binding residues. This treble-clef core is elaborated by distinct surrounding secondary-structure elements, including, in one case (EPP00000606), a C-terminal transmembrane helix (rightmost structure in **Fig 7A**).

**Figure 7.**
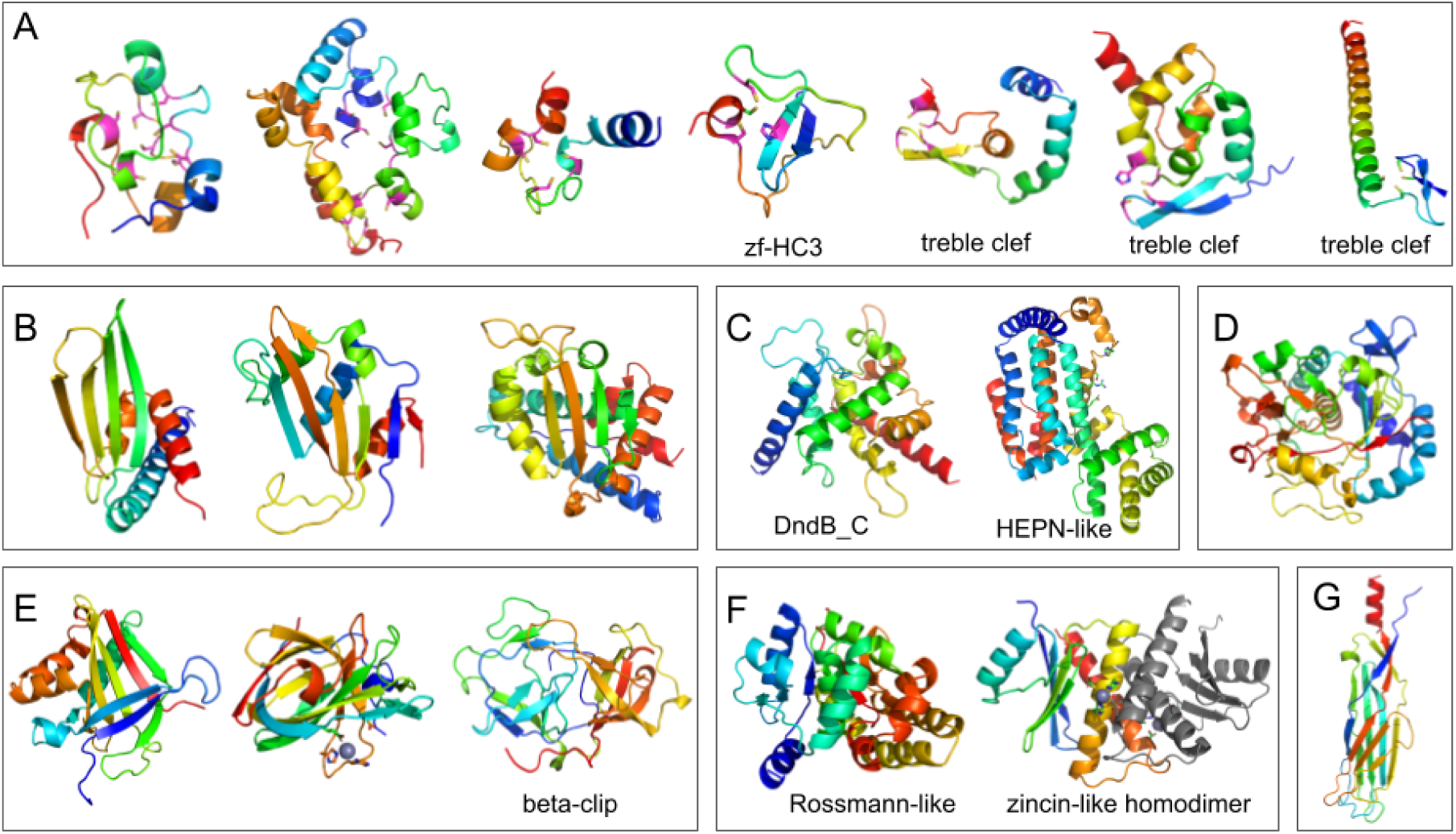
Examples of AlphaFold structural models of genuinely dark domains. **(A)** Cysteine-rich metal-binding domains. Conserved cysteine and histidine residues that likely coordinate metal ions are shown in pink. **(B)** Three a+b two-layered domains with potential novel folds. **(C)** Two alpha-helical domains likely involved in defense. **(D)** A domain with complex topology representing a potential novel fold. **(E)** Three beta-barrel domains. **(F)** Two a+b two-layered domains. The right structure adopts a Rossmann-like fold. The left structure is modeled as a homodimer with two zinc ions using AlphaFold3; one subunit is colored in a rainbow scheme and the other in gray. Side chains of conserved metal-binding and catalytic residues are shown as sticks. **(G)** A beta-sandwich domain with a potential novel fold.

Other architectures include a+b two-layer folds (a single b-sheet with a-helices on one side; 6 cases), a-helical arrays or bundles (5 cases), beta-barrels (2 cases), a+b three-layer folds (2 cases), beta-sandwiches (2 cases), and a+b complex topology (2 cases). Several domains appear to represent novel folds, including three a+b two-layered domains (**Fig 7C**, A0A060HJU8, EPP00026547 and EPP00002661), one a+b domain with complex topology (**Fig 7D**, EPP00003247), two beta-barrels (**Fig 7E**, left and middle structures, accession: EPP00010451 and EPP00001136), and one beta-sandwich domain (**Fig 7G**, EPP00002877).

Detailed manual analysis of sensitive sequence and structural similarity searches further suggests that some of these domains may be remote homologs of known protein families. For example, one alpha-helical domain (accession: EPP00023006) was identified as a distant homolog of the C-terminal region of DndB (DndB_C; **Fig 7C**, left structure, HHpred score: 98.51), which is involved in DNA phosphorothioate modification [28]. Another alpha-helical bundle domain (EPP00023009) shows similarity to the HEPN superfamily of endoribonucleases [29] and contains several conserved residues that are likely to form the active site (**Fig 7D**). In addition, one case (UPI00355E9374) represents a case of duplication of beta-clip fold domains, which are found in prokaryotic phage capsid decoration (cement) proteins located on the outside surface of the icosahedral capsid [30].

We also identified a domain with a Rossmann-like fold (**Fig 7F**, left structure, UPI003165088D) that does not exhibit detectable sequence similarity to any known domains. Structural searches using DaliLite identified domains in the HAD superfamily of phospho-hydrolases [31] as the closest matches, characterized by short beta-strands and two alpha-helices in the Rossmann-fold crossover region. However, this domain lacks the canonical metal-binding active-site residues of HAD enzymes and contains additional C-terminal secondary-structure elements, including an antiparallel beta-strand. Thus, its potential evolutionary relationship to HAD domains remains unclear.

Another a+b three-layered protein domain (**Fig 7F**, right structure, UPI00355F993C) shows structural similarity to zincin domains and contains the characteristic HEXXH motif of zincin-like metalloproteases [32]. However, in monomeric models, the third conserved acidic residue located in the final alpha-helix is positioned distal to the HEXXH motif, unlike in canonical zincins. When modeled as a homodimer (one subunit colored rainbow and the other colored gray in **Fig 7F** right structure) using AlphaFold3 in the presence of two zinc ions, the acidic residue from the adjacent monomer is positioned close to the HEXXH motif, completing the metal-binding site at the dimer interface. To our knowledge, this represents a unique arrangement among zincin-like metalloproteases.

### MCR and MVP illustrate lineage-specific and pan-archaeal domain distributions

To illustrate the biological information captured by pan-archaeal domain classification, we highlight two protein families with contrasting phylogenetic distributions (**Fig 8**).

**Figure 8.**
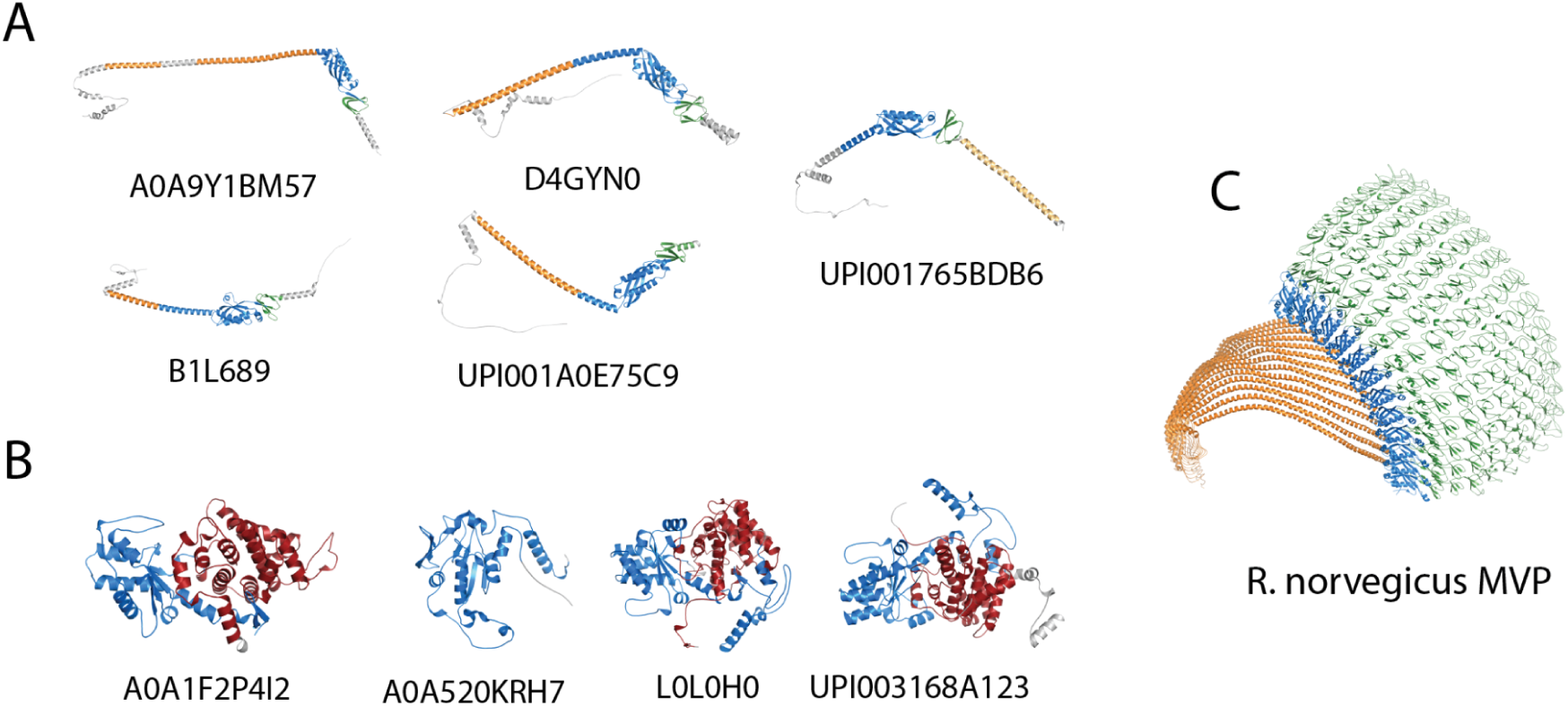
Case study structures illustrating lineage-specific and pan-archaeal domain distributions. **(A)** Methyl-coenzyme M reductase (MCR) representatives from functionally diverse lineages, colored by ECOD domain: blue = H-group 304.35 (MCR N-terminal), red = H-group 154.1 (MCR C-terminal), gray = unassigned. The conserved two-domain architecture is maintained across a Syntrophoarchaeum alkane oxidizer (A0A1F2P4I2, 460 aa), deep-branching Hadarchaeales (UPI003168A123, 607 aa), and a TACK methylotroph (L0L0H0, 571 aa), with a single-domain variant from Methanoliparum (A0A520KRH7, 262 aa). MCR domains are archaea-exclusive (zero bacterial, zero eukaryotic in ECOD) and distributed by metabolism rather than phylogeny. **(B)** Major vault protein (MVP) representatives from all five major archaeal groups, colored by ECOD domain: green = H-group 3529.1 (MVP repeat), blue = H-group 230.5 (SPFH/Band 7), orange = lineage-variable C-terminal domains, gray = unassigned. The conserved repeat-shoulder-tail architecture spans Asgard (A0A9Y1BM57, 392 aa), DPANN (A0A842X616, 322 aa), Euryarchaeota (D4GYN0, 396 aa), TACK (B1L689, 328 aa), and deep-branching archaea (A0A7C1HSU6, 317 aa). **(C)** Eukaryotic MVP from rat liver vault (PDB 2ZUO), shown as a single chain (845 aa) and as the 13-chain half-vault assembly, with the same color scheme. The eukaryotic monomer elaborates the archaeal architecture with 8 tandem MVP repeats (green), a retained SPFH shoulder (blue), and a helical cap domain (orange, H-group 3530.1).

Methyl-coenzyme M reductase (MCR) exemplifies a lineage-restricted, archaea-exclusive enzyme family. The two MCR domains (H-groups 304.35 and 154.1) have zero bacterial and zero eukaryotic representatives in ECOD. Within our dataset, MCR domains are present in classical methanogens (*Methanosarcina*, *Methanococcus*, *Methanopyrus*), but also in organisms that use MCR-like enzymes for anaerobic alkane oxidation rather than methanogenesis: including *Ca. Syntrophoarchaeum butanivorans* (anaerobic butane oxidizer; 20 MCR domains, the most in our dataset) and *Ca. Hadarchaeales* (deep-branching archaea with proposed hydrocarbon metabolism). Bathyarchaeota carry MCR homologs whose function remains debated. The conserved two-domain architecture, an N-terminal alpha/beta domain (304.35) packed against a C-terminal helical domain (154.1), is maintained across these functionally diverse lineages, with size variation (262–607 aa) but not domain organization (**Fig 8B**). MCR domains are absent from Asgard, DPANN, and non-methanogenic Euryarchaeota, consistent with a metabolism-linked rather than phylogeny-linked distribution. Domain classification identifies the shared structural scaffold but cannot distinguish the reaction direction; the same two-domain architecture supports both methane production and alkane activation.

The major vault protein (MVP) illustrates the opposite pattern. MVP was recently reported as an Asgard-eukaryote structural connection by Kostlbacher et al. [24], who identified Asgard proteins with reciprocal best structural hits to eukaryotic MVP. However, the MVP structural repeat (H-group 3529.1) has been classified in ECOD since 2014 (PDB 2ZUO, rat liver vault). Our pan-archaeal classification identifies 32 high-confidence MVP domains across all six major archaeal groups, with Asgard accounting for only 5 (16%). The conserved multi-domain architecture; an N-terminal MVP repeat (3529.1), a central SPFH/Band 7 domain (230.5), and lineage-variable C-terminal domains, is present from DPANN to Asgard (**Fig 8A**). In the eukaryotic vault (2ZUO), the same architecture is elaborated: 8 tandem MVP repeats replace the single archaeal repeat, the SPFH shoulder is retained, and a long helical cap domain (3530.1) extends into the vault particle interior (**Fig 8C**). The archaeal proteins thus represent a compact version of the eukaryotic vault monomer. Whether archaeal MVPs assemble into vault-like particles is unknown, but the domain-level conservation suggests that the vault architecture predates the archaeal-eukaryotic divergence rather than originating as an Asgard-specific innovation. The distinction between Asgard-specific and pan-archaeal distribution is only visible with the breadth of sampling that pan-archaeal classification provides.

### The archaeal fold repertoire in context

Combining direct classification with structural clustering rescue, the accounting of the 124,075 archaeal proteins is as follows. A total of 98,110 proteins (79.1%) are classified directly by DPAM with high confidence, and an additional 3,463 (2.8%) are reclassifiable through structural cluster transitivity. The sub-threshold fraction, 17,513 (14.1%) proteins with domain matches below the confidence threshold, represent known homologous groups detected at reduced sensitivity, not structural novelty. Among the genuinely domain-free proteins, the majority are explained by low-confidence structure proteins (5.0%), short length below classification sensitivity (0.9%), or lack of independent structural support (0.1%). A small number of candidates for genuine structural novelty remain under investigation. Existing homologous groups thus account for virtually all confidently predicted archaeal protein structures. The sub-threshold fraction, the single largest component of the unclassified proteome, represents a classification sensitivity gap rather than a structural novelty gap: these proteins have detectable similarity to known folds, just below the threshold for high-confidence (and automated) assignment.

## Discussion

The central finding of this study is that the protein fold repertoire at the single-domain level is broadly conserved across cellular life. Archaea, the most phylogenetically distant and structurally undersampled domain, are built from the same set of domain-level building blocks as bacteria and eukaryotes. Of 124,075 proteins spanning all major archaeal lineages, approximately 80% receive high-confidence domain classifications directly, and systematic characterization of the unclassified fraction reveals that it is dominated by classification sensitivity limits and structure prediction quality rather than by novel structures.

### The unclassified fraction represents sensitivity, not novelty

The elevated orphan fraction in archaeal domains (18.2% versus 7.7% in Swiss-Prot) initially suggests extensive uncharacterized structural diversity. However, our analysis reveals that this gap is driven primarily by classification sensitivity. The largest contributor is structure prediction quality: 74% of proteins lacking any domain assignment have mean pLDDT scores below 70, indicating disordered regions or unreliable predictions rather than well-folded domains of unknown type. Among well-folded, reasonably sized domain-free proteins, structural clustering connects most to proteins that do carry DPAM classifications, either directly (classifiable by transitivity) or through sub-threshold matches (partial signal). The funnel from 8,452 domain-free proteins to a small number of genuinely uncharacterized candidates reflects a series of straightforward explanations, poor predictions, short sequences, known folds at sub-threshold confidence, that collectively account for the majority of the unclassified fraction.

The 14% sub-threshold fraction merits particular attention. These proteins have DPAM domain matches to known folds, just below the confidence threshold used for high-confidence classification. Structural clustering validates this interpretation: in mixed clusters containing both high-confidence and sub-threshold members, 81.5% of sub-threshold domains carry X-group assignments that agree with the cluster consensus. The dominant issue is calibration, the sensitivity of template-based classification for deeply divergent archaeal sequences, not the absence of matching templates. This finding suggests that the next gains in archaeal classification will come from improving classifier sensitivity rather than from discovering new folds.

### Conservation of the topology repertoire across cellular life

The question of whether the protein fold universe is finite and mappable has recurred across multiple eras of structural biology. Structural genomics initiatives of the early 2000s consistently found that new folds appeared at a diminishing rate [33–35], and estimates of the total number of domain-level folds converged on a range of 1,000 to 10,000 [36–38]. The predicted structure revolution has revisited this question at a qualitatively different scale: where structural genomics solved thousands of structures per year, AlphaFold now provides models for hundreds of millions of proteins. At each stage of expansion, from PDB-only to predicted structures from 48 model organisms [3, 26] to the archaeal proteomes surveyed here, the proportion of genuinely novel folds discovered has been small relative to the expansion of coverage within known fold families.

Our results extend this pattern to the most phylogenetically extreme test available within cellular organisms. Archaea include lineages (Asgardarchaeota, DPANN, deep-branching thermophiles) with no close relatives among well-characterized organisms. If large regions of fold space were accessible only through deeply divergent lineages, archaeal proteomes should reveal them. That existing templates instead cover archaeal proteins at rates comparable to bacterial and eukaryotic proteomes suggests that the domain-level fold repertoire is shared across cellular life, a statement about the universality of the structural building blocks, not an enumeration of all possible folds.

Several important caveats apply. First, this conclusion is restricted to the single-domain level; the combinatorial space of domain arrangements is far larger than the space of individual folds and remains poorly characterized. Second, and critically, structure prediction methods are inherently conservative: AlphaFold is trained on known structures and produces confident predictions most readily for proteins that resemble its training data. The pLDDT filter that removes 5% of proteins as “disordered or poorly predicted” may also exclude proteins whose structures are genuinely novel but fall outside the prediction model’s confident regime. We cannot fully distinguish between “no novel folds exist” and “our methods cannot confidently predict novel folds.” The strongest version of our claim is comparative: archaeal proteins are classified at the same rate as other cellular proteomes, indicating that archaea are not a reservoir of unexplored fold diversity, but this does not preclude the existence of novel folds among proteins for which no confident prediction is available. A systematic test of whether X-group boundaries, which represent uncertain evolutionary relationships between homologous superfamilies, are robust to archaeal diversity found no new cross-X-group relationships among archaeal domains, consistent with the overall conservation finding.

### Data provenance and confidence

The phylogenetic breadth that makes this dataset a strong test of fold repertoire conservation is itself dependent on metagenome-assembled genomes. Of the 65 GTDB classes in our dataset, approximately 60% are represented primarily or exclusively by MAG-derived organisms, including the Asgard archaea, much of DPANN, and most deep-branching lineages. Without metagenomics, systematic structural coverage of archaeal diversity would be limited to the ∼18 cultivated classes.

The three data sources carry different levels of confidence. Proteins from the AlphaFold Database (39 classes, 58% of the dataset) have curated gene models and structures validated through the AFDB pipeline. UniParc-source proteins (11 classes, 18%) have established sequences but require de novo structure prediction. Prodigal-source proteins (15 classes, 24%) carry compound uncertainty: gene boundaries, protein sequences, and structures are all derived computationally from metagenomic assemblies of varying quality. This provenance gradient is visible in the dark matter analysis, where Prodigal-source proteins are overrepresented among uncharacterized candidates relative to their share of the full dataset, consistent with gene prediction artifacts inflating the apparent novelty. The conservation finding is robust to data source: even restricting to AFDB-source proteins alone, the fraction of well-folded structural orphans remains below 0.1%.

### Open frontiers

These results point to several productive directions for structural classification. The most immediate is family-level expansion. The ∼1,000 fold-level building blocks may be broadly shared across cellular life, but family-level membership, the sequence and structural diversity within each fold, is where lineage-specific adaptation lives. Comprehensive classification of the full predicted structure universe, now encompassing hundreds of millions of proteins, is the natural next step for mapping this diversity.

Viral proteomes represent a qualitatively different frontier. Viruses evolve under fundamentally different constraints from cellular organisms, including de novo gene origination and capsid architecture, and are not bound by the same evolutionary continuity that links archaeal, bacterial, and eukaryotic folds. If genuinely novel folds exist beyond the cellular repertoire, viral proteomes are the most promising place to look.

Multi-domain architectures constitute another open question. Our analysis is restricted to the single-domain level, but the combinatorial space of domain arrangements is far larger than the space of individual folds. Domain co-occurrence rules, architecture evolution, and lineage-specific domain combinations remain poorly characterized and represent a substantial source of protein functional diversity.

Finally, classifier sensitivity is a tractable engineering problem. The 14% sub-threshold fraction, proteins with detectable but sub-threshold similarity to known folds, represents the largest opportunity for improving archaeal classification coverage. The high-confidence threshold used for automated ECOD accession reflects an operational trade-off between classification accuracy and throughput, and many sub-threshold matches are likely correct but fall below the confidence level at which assignments can be made without manual review. Structural clustering offers one path to promoting these assignments, our cluster-based validation suggests that most sub-threshold domains are correctly assigned at the X-group level, but using structural similarity as a direct proxy for evolutionary classification carries its own risks, particularly for superfolds where convergent evolution can place unrelated domains in the same structural cluster. The path forward likely involves combining expanded template libraries, improved profile-based search for divergent sequences, and cluster-informed classification with appropriate caution about the distinction between structural similarity and homology.

## Conclusions

Systematic structural classification of archaeal proteomes shows that the protein fold repertoire at the single-domain level is broadly conserved across all major cellular lineages. The gap between archaeal and well-characterized proteomes reflects classification sensitivity for divergent sequences, not an abundance of novel structures. The priorities for structural classification accordingly shift from fold discovery toward family-level expansion, sensitivity improvements, and characterization of the combinatorial diversity of domain architectures that these conserved building blocks assemble into.

## Methods

### Genome selection and protein dataset assembly

Archaeal genomes were selected from the Genome Taxonomy Database (GTDB) release r220 [39], which contains 11,918 archaeal genome assemblies (1,178 cultivated and 10,740 metagenome-assembled genomes [MAGs]). To complement existing AlphaFold Database (AFDB) coverage, we excluded genomes already represented in AFDB (8,800 MAGs encompassing ∼14.3 million proteins) and targeted the remaining genomes for download from NCBI. Protein sequences were obtained for 954 genomes: 367 cultivated genomes (123 from UniProt reference proteomes, 244 from NCBI GenBank/RefSeq) and 587 MAGs. MAGs were classified by CheckM quality metrics: 236 as high-quality (completeness ≥90%, contamination ≤5%) and 351 as medium-quality (completeness ≥50%, contamination ≤10%).

From this pool, a representative working set of 65 GTDB classes was curated to balance phylogenetic coverage across known archaeal lineages, inclusion of both cultivated organisms and MAGs, and representation of phylogenetically distinct groups that are undersampled in existing structural databases. These 65 classes span 21 phyla and were organized into 6 operational groups based on established archaeal phylogeny: Asgard, DPANN, TACK superphylum, Euryarchaeota-related, Deep-branching, and Thermoplasmatales. The working set comprises 124,075 proteins drawn from three sources that differ in provenance and confidence. The first source (39 classes, 71,866 proteins) consists of proteins with existing AlphaFold2 structures in the AFDB, representing a mix of cultivated reference organisms with curated gene models and MAG-derived organisms already accessioned by AFDB. The second source (11 classes, 22,883 proteins) consists of proteins with existing sequences in UniParc but no AFDB structures; for these, we predicted structures de novo using AlphaFold3. The third source (15 classes, 29,326 proteins) consists of proteins from lineages with neither AFDB structures nor UniParc sequences; for these, we performed gene prediction with Prodigal [40] on MAG assemblies and then predicted structures with AlphaFold3. This third category carries compound uncertainty: gene boundaries, protein sequences, and predicted structures are all derived computationally from metagenomic assemblies, but includes many phylogenetically important lineages (e.g., Baldrarchaeia, Hermodarchaeia, Wukongarchaeia) that are not otherwise represented in structural databases.

### Structure prediction and quality assessment

Existing AlphaFold2 structures for the AFDB subset were retrieved from AFDB version 6. For the remaining 52,209 proteins (Prodigal and UniParc sources), multiple sequence alignments were generated using HHblits 3.3.0 [41] with 3 iterations against the UniRef30 2023_02 database [42] (E-value threshold 0.001). Structures were then predicted using AlphaFold3 v1 [18] with pre-computed MSAs (--norun_data_pipeline) on the TACC Lonestar6 computing cluster (NVIDIA A100 GPUs). Proteins were partitioned into tiers by length (Tier 0.5: 50–100 aa; Tier 1: 100–500 aa; Tier 2: 500–1000 aa; Tier 3: >1000 aa) and processed in batches of 6 parallel GPU tasks per SLURM job. AF3 outputs were converted to AFDB-compatible mmCIF and PAE JSON formats for downstream processing.

Per-protein structure quality metrics were extracted from predicted structures for 116,275 proteins (93.7% of the dataset). Metrics included mean per-residue pLDDT confidence score (extracted from B-factor fields), fraction of disordered residues (pLDDT < 50), predicted TM-score, secondary structure composition (helix, sheet, and coil fractions), radius of gyration, and an overall quality category. The mean pLDDT score served as the primary indicator of structure prediction confidence and was used as a filter in the dark matter analysis.

### Domain assignment and annotation

Protein domains were identified using the Domain Parser for AlphaFold Models (DPAM)[19], which assigns ECOD[3] domain boundaries and classifications through iterative profile-profile search (HHsearch)[41], structural comparison (Foldseek, Dali)[22, 43], domain boundary refinement, and confidence scoring. DPAM was run against ECOD version 292 (June 2025). DPAM’s primary classification output is a T-group (topology group) assignment, from which higher-level groupings, H-group (homologous superfamily) and X-group (possible homology), are derived by traversing the ECOD hierarchy. Each domain also receives a classification judge reflecting confidence: “good domain” (high-confidence ECOD match), “simple topology” (matched to a common fold with low specificity), “low confidence” (weak structural match), or “partial domain” (incomplete boundary). The “good domain” classification was used as the primary quality filter in downstream analyses.

DPAM was run on the LEDA HPC cluster using SLURM array parallelization. Separate processing runs accommodated the different input sources (AFDB structures, de novo AF3 predictions, and proteins requiring additional profile searches), but all used the same DPAM pipeline and reference data. Domain sequences were extracted from parent protein sequences using DPAM-assigned residue ranges.

Domain sequences were further annotated against Pfam-A version 38.2 using hmmscan from HMMER 3.1b2 [44] with the Pfam gathering threshold (GA) cutoff. Results were parsed from HMMER domain table output, extracting per-domain E-values, alignment coordinates, and posterior probabilities. In the standard ECOD accession workflow, T-group assignments from DPAM are combined with Pfam analysis to determine F-group (family) assignments [26].

### Sequence and structural clustering

Clustering was performed at four levels to enable systematic comparison of sequence-based and structure-based groupings. Domain-level clustering used sequences and structures extracted at DPAM-assigned boundaries, while protein-level clustering used full-length sequences and structures.

#### Protein sequence clusters (PSC)

Full-length protein sequences were clustered with MMseqs2 [45] at 40% minimum sequence identity with 80% bidirectional coverage (`--min-seq-id 0.4 -c 0.8 --cov-mode 0`).

#### Domain sequence clusters (DSC)

Domain sequences were clustered with MMseqs2 using the same parameters as PSC (`--min-seq-id 0.4 -c 0.8 --cov-mode 0`).

#### Protein structural clusters (PXC)

Full-length protein structures were clustered with Foldseek (van Kempen et al., 2024) using a three-step pipeline: database creation (`createdb`), all-vs-all search, and greedy set-cover clustering (`clust --cluster-mode 0`). Search parameters were: TM-align alignment mode (`--alignment-type 2`), E-value threshold 0.001, minimum coverage 0.5, and no sequence identity filter (`--min-seq-id 0.0`).

#### Domain structural clusters (DXC)

Domain structures (extracted as individual mmCIF files) were clustered with Foldseek using the same pipeline and parameters as PXC.

Structural clustering parameters (E-value 0.001, 50% coverage, TM-align mode) were chosen to capture remote structural relationships while filtering spurious matches; TM-align mode scores full structural superposition rather than local fragment matching, appropriate for the domain-level comparisons central to this study.

### Cross-clustering analysis

To quantify the extent to which structural clustering reveals relationships invisible to sequence comparison, we computed cross-cluster mappings between structural and sequence cluster dimensions. For each structural cluster, we identified the set of sequence clusters represented by its members. This analysis was performed at both protein (PXC to PSC) and domain (DXC to DSC) levels.

### Characterization of the unclassified fraction

Proteins lacking any high-confidence DPAM domain assignment were systematically characterized through a series of filters designed to distinguish genuine structural novelty from explainable non-classification.

Proteins were first partitioned by DPAM classification status: “classified” (at least one domain with the “good domain” judge), “sub-threshold” (at least one DPAM domain of any judge category, but none reaching “good domain”), and “no domain” (no DPAM domain assignments at all). The no-domain category was further filtered to identify well-folded candidates for structural novelty. Proteins with a mean pLDDT below 70 were excluded as disordered or poorly predicted structures, and proteins shorter than 100 amino acids were excluded as below the effective sensitivity of template-based domain classification. The remaining proteins were assessed through their PXC cluster memberships: singletons (lacking any structural neighbor) were excluded, and proteins in clusters containing at least one member with a high-confidence domain assignment were classified as identifiable by transitivity. Proteins in clusters where at least one member carried any DPAM domain assignment (even sub-threshold) were classified as having partial signal. The residual, proteins in fully dark clusters where no member has any DPAM domain assignment at any confidence level, constitutes the irreducible core of structurally uncharacterized proteins.

### Domain rescue by structural clustering

To assess whether structural clustering could validate sub-threshold DPAM classifications, we identified DXC clusters containing both high-confidence (“good domain”) and sub-threshold members. For each such mixed cluster, we determined the X-group consensus among the high-confidence members. Clusters where all high-confidence members shared the same X-group were designated as having consistent consensus. The X-group assignments of sub-threshold domains in consistent clusters were then compared against the cluster consensus and scored as agreeing, disagreeing, or receiving a new assignment (for domains lacking a prior X-group assignment). Protein-level rescue was defined as upgrading a protein from unclassified to classifiable when at least one of its sub-threshold domains was validated by a consistent rescue cluster.

### Comparative analyses

Archaeal domain statistics were compared against our classification of Swiss-Prot[26] to contextualize classification coverage. Swiss-Prot protein lengths, domain classifications, and Pfam mappings were obtained from the UniProt and ECOD databases. ECOD hierarchy totals (X-groups, H-groups, T-groups) were computed from the full ECOD v292 classification.

### Data and code availability

All analysis scripts, figure-generation code, and frozen data exports are available at Zenodo (DOI: 10.5281/zenodo.19269332). The repository includes domain assignments for 204,758 domains across 124,075 archaeal proteins, protein- and domain-level sequence and structure clustering results, Pfam annotations, and all data required to reproduce manuscript figures without database access. Figure scripts support both database-connected and CSV-only execution modes for full reproducibility. Predicted structures and domain assignments are additionally available at http://prodata.swmed.edu/ecod2/epp.

### Use of AI-assisted tools

Claude Opus 4.6 and Sonnet 4.6 (Anthropic) were used as a programming assistant during data analysis, figure generation, and manuscript preparation. Additionally, ChatGPT 5.3 was used for some manuscript preparation. All AI-assisted code was reviewed and validated by the authors. Claude was used for: database query construction and optimization, Python scripting for figure generation and data analysis, literature search assistance, and manuscript copyediting. All scientific interpretations, experimental design decisions, and domain expertise judgments were made by the authors. The AI tool did not generate scientific hypotheses, design experiments, or make classification decisions.

## Notes

### Competing Interest Statement

The authors have declared no competing interest.

